# EYE-Llama, an in-domain large language model for ophthalmology

**DOI:** 10.1101/2024.04.26.591355

**Authors:** Tania Haghighi, Sina Gholami, Jared Todd Sokol, Enaika Kishnani, Adnan Ahsaniyan, Holakou Rahmanian, Fares Hedayati, Theodore Leng, Minhaj Nur Alam

## Abstract

1

**Background:** Training Large Language Models (LLMs) with in-domain data can significantly enhance their performance, leading to more accurate and reliable question-answering (Q&A) systems essential for supporting clinical decision-making and educating patients.

**Methods:** This study introduces ophthalmic LLMs trained on in-domain, well-curated datasets. We present an open-source substantial ophthalmic language dataset for model training. Our models (EYE-Llama), were pre-trained on an ophthalmology-specific dataset, including paper abstracts, textbooks, and Wikipedia articles. Subsequently, the models underwent fine-tuning using a diverse range of QA pairs. Our models were compared to baseline Llama 2, ChatDoctor, Meditron, Llama 3, and ChatGPT (GPT3·5) models, using four distinct test sets, and evaluated quantitatively (Accuracy, F1 score, BERTScore, BARTScore and BLEU score) and qualitatively by two ophthalmologists.

**Findings:** Upon evaluating the models using the synthetic dialogue test set with three different metrics (BERTScore, BARTScore, and BLEU score), our models demonstrated superior performance. Specifically, when evaluated using BERTScore, our models surpassed Llama 2, Llama 3, Meditron, and ChatDoctor in terms of F1 score, and performed on par with ChatGPT, which was trained with 175 billion parameters (EYE-Llama: 0.57, Llama 2: 0.56, Llama 3: 0.55, Meditron: 0.50, ChatDoctor: 0.56, and ChatGPT: 0.57). Additionally, the EYE-Llama model outperformed the above models when evaluated using BARTScore and BLEU scores. When tested on the MedMCQA test set, the fine-tuned models exhibited higher accuracy compared to Llama 2, Meditron, and ChatDoctor models (EYE-Llama: 0.39, Llama 2: 0.33, ChatDoctor: 0.29, Meditron: 0.22). However, ChatGPT, and Llama 3 models outperformed EYE-Llama, achieving accuracies of 0.55, 0.78, and 0.90, respectively. On the PubmedQA test set, our model showed improved accuracy over all other models (EYE-Llama: 0.96, Llama 2: 0.90, Llama 3: 0.92, Meditron: 0.76, ChatGPT: 0.93, ChatDoctor: 0.92).

**Interpretation:** The study shows that pre-training and fine-tuning LLMs like EYE-Llama enhances their performance in specific medical domains. Our EYE-Llama models surpass baseline Llama 2 in all evaluations, highlighting the effectiveness of specialized LLMs in medical QA systems.

**Funding:** Funded by NEI R15EY035804 (MNA), R21EY035271 (MNA), and UNC Charlotte Faculty Research Grant (MNA)

## 2. Introduction

Generative large-language models (LLMs) have transformed Natural Language Processing (NLP), creating human-sounding text and scoring superbly across a broad range of linguistic tasks. Clinically, they can transform the workload of clinicians by taking over routine documentation and triage, freeing up professionals to spend time on direct patient care. Chatbots and decision aids made with LLMs can also provide personalized treatment and promote participation by delivering individualized education and self-monitoring tools. With these advantages, LLM-based systems will soon become staples of contemporary medicine. Two impediments continue to retard large-scale adoption: an insufficiency of high-quality, domain-specific data for fine-tuning, and the prohibitive computational expense of training and serving extremely large models.

Despite these hurdles, popularity for medical LLMs has developed exponentially in the past few years.^1–3^ Most of today’s work compares in-off-the-shelf models to clinical databases, assessing their ability to answer multiple-choice (MC) and open-ended questions.^4,5^ On both types, ChatGPT is one of the high standards set by general-purpose models.^6^ Llama 2 is equally as accurate with many fewer parameters, though.^7^ The latest iteration of Llama 3 again raises the ceiling, with superior performance on question-answering tasks.^8^ More critically, each of these models further narrows the gap when fine-tuned on in-domain corpora. Examples such as ChatDoctor, BioGPT, Me-LLaMA, and PMC-LLaMA illustrate this strategy by fine-tuning general models on medical text to improve clinical Q&A performance.^9–12^ Similarly, Meditron, pre-trained on biomedical literature alone, has performed well on specialty benchmarks.^13^ However, additional tuning is needed before such systems can return reliably accurate answers in subspecialties such as ophthalmology.

The contributions of our study are as follows:

- We have developed and open-sourced a comprehensive dataset in ophthalmology. The data is composed of roughly 744k samples collected from PubMed abstracts, 22k samples from nearly 570 textbooks, and online articles. Additionally, our dataset includes supervised data, comprising approximately 18k Q&A pairs from medical datasets, 1·5k synthetic dialogues, as well as 15k Q&A pairs generated using GPT3·5.
- We have proposed a framework for developing an open-source model that was trained on in-domain datasets and tailored to specific areas within medicine, such as ophthalmology. (*DOI: 10.5281/zenodo.15272005*).
- We have investigated and validated our model against baseline LLMs, to prove the feasibility of our hypothesis, that is: the in-domain dataset enhances LLM performance, even in smaller LLMs with reduced trainable parameters.

## 3. Methods

Figure 1 outlines the training workflow. We adopt a two-stage strategy to adapt Llama 2-7B-Chat for ophthalmology. First, we pre-train the base model on a broad corpus of domain-specific texts, producing the intermediate checkpoint **EYE-Llama_p**. In the second stage, we fine-tune that checkpoint on carefully assembled ophthalmic Q&A sets. The smaller collection yields **EYE-Llama_qa**, while an expanded version, augmented with an additional 15,000 Q&As, produces **EYE-Llama_gqa**. This progression lets the model absorb general ophthalmic language before specializing in detailed clinical questioning.

**Figure 1.**
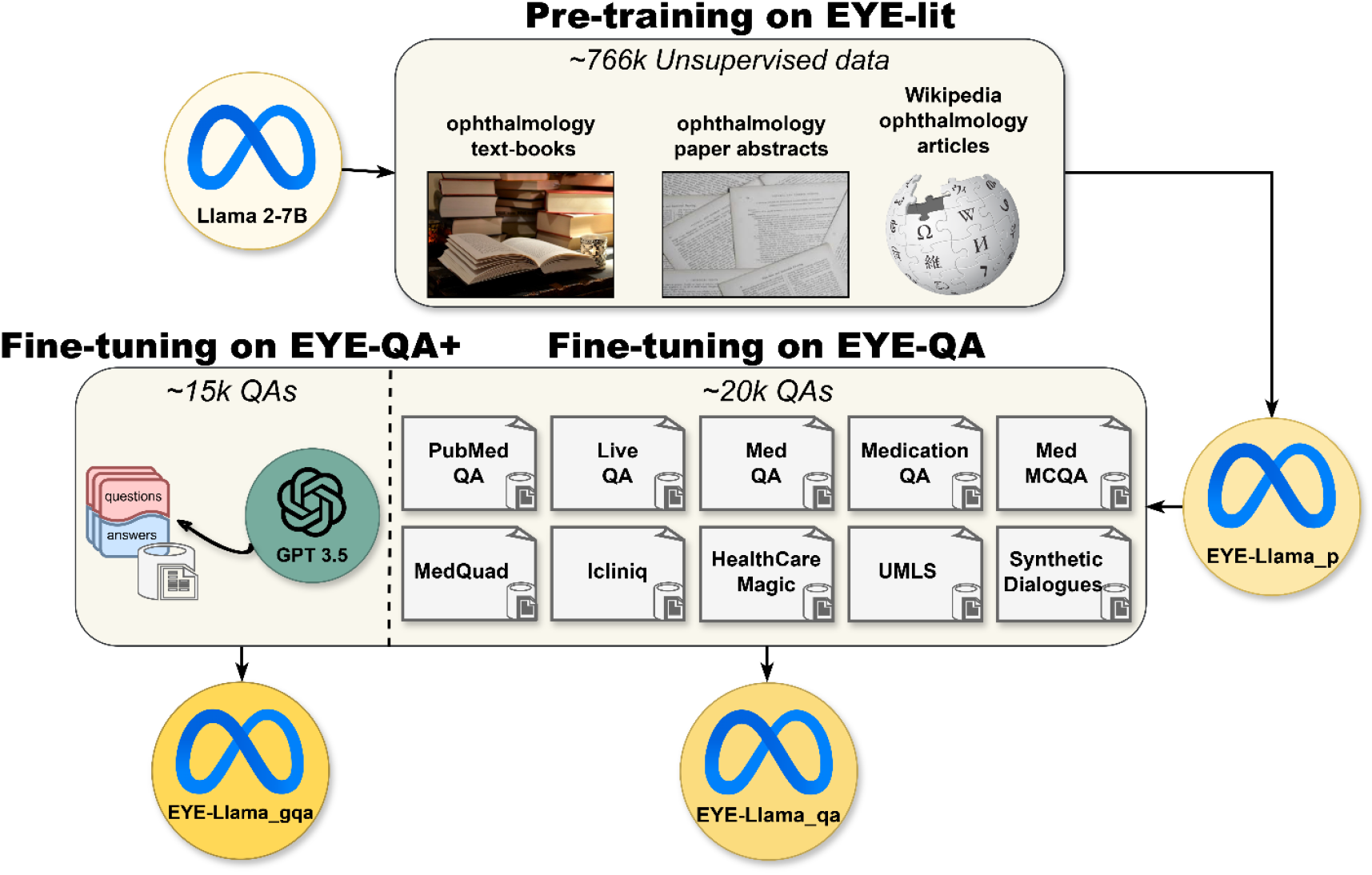
Schematic representation of the Llama 2 model’s pre-training and fine-tuning process.

### 3.1 Dataset

#### 3.1.1 Unsupervised data

We began by harvesting roughly 744,000 ophthalmic abstracts from PubMed. Using BioPython’s Entrez package, we issued keyword-based queries to the NCBI database and downloaded every abstract that contained at least one ophthalmology-related term. The search terms were: *<u>ophthalmology, eye, conjunctiva, retina, sclera, cone cell, choroid, macula, cataract, strabismus, optic nerve, macular degeneration, ciliary body, ophthalmologist, pinkeye, astigmatism, diabetic eye disease, diabetic retinopathy, age-related macular degeneration, glaucoma, iris, cornea, optometry, and geographic atrophy</u>*. These same terms later served as filters for the downstream Q&A datasets shown in Figure 1 (e.g., PubMedQA, MedMCQA).

To broaden topic coverage and stylistic variety, we supplemented the abstracts with two additional sources: 570 ophthalmology textbooks and full-length Wikipedia articles drawn from the platform’s ophthalmology category (see Appendix C). Collectively, this corpus provides the linguistic and factual breadth needed to pre-train an LLM for eye-care applications.

**Table 1.** EYE-lit dataset description.

| EYE-lit Dataset |  |  |  |
| --- | --- | --- | --- |
| Source | Percentage | License | Number of Tokens |
| Online articles | 2% | Open access | ~5m |
| PubMed Abstracts | 98% | Open access | ~248m |

**EYE-lit** is purpose-built for ophthalmology: rather than mixing in general medical texts, it contains only eye-care literature. This strict focus on domain terminology and concepts gives language models the detailed exposure they need to master the field’s distinctive vocabulary and nuances (see Appendix F).

#### 3.1.2 Data cleaning

We first converted the PubMed abstracts from XML to plain text, stripping out special characters and boiler-plate phrases such as “in this paper”. Textbook chapters and Wikipedia articles were then sliced into self-contained chunks of no more than 512 tokens. Because nearly all PubMed abstracts have already fallen below this threshold, we left them intact and segmented the other sources to match, ensuring consistent paragraph length across the corpus. The resulting collection, now dubbed EYE-lit, contains only complete, information-rich passages. It serves as the foundation for our pre-training experiments and is publicly available at huggingface.co/datasets/QIAIUNCC/EYE-lit-complete.

#### 3.1.3 Supervised data

Supervised fine-tuning began with a carefully chosen slice of the large medical Q&A collection published by Chaoyi Wu et al.^11^ Using the same ophthalmology keywords that guided our PubMed search, we pulled matching Q&A pairs from that corpus, the Icliniq and HealthCareMagic forums, and the MedQuad set.^9,14^ The harvest came to roughly 18,000 items: about 6,000 each from PubMedQA and MedMCQA for training, plus another 800 from each for testing.^15,16^

Because real-world dialogue rarely reads like an MC exam, we created an additional set of patient-style Q&As. GPT3.5 played the role of an ophthalmologist, answering queries that our clinicians later checked for plausibility. One prompt (Appendix A1) yielded about 2800 exchanges; we kept 1500 for training and 1300 for evaluation (available at huggingface.co/datasets/QIAIUNCC/EYE-Synthetic-Dialogues).

Next, we folded everything into EYE-QA, our unified ophthalmic Q&A corpus. To help the model cope with more specialized prompts, say the sort of quick consultation an ophthalmologist might throw at an AI assistant, we generated roughly 15,000 extra textbook-based Q&As with GPT3.5 Turbo (prompt in Appendix A2). Adding these to EYE-QA produced EYE-QA+, which is available at huggingface.co/datasets/QIAIUNCC/EYE-QA-PLUS.

EYE-QA+ dataset:

- Subset of medical QA datasets (18k samples)
  ∘ PMC_LLaMA_instructions dataset
  ∘ Icliniq and HealthCareMagic datasets
  ∘ MedQuad dataset
- Patient/ophthalmologist synthetic dialogues generated with GPT3.5 (1·5k)
- Synthetic QA pairs generated by GPT3.5 (15k)

**Figure 2.**
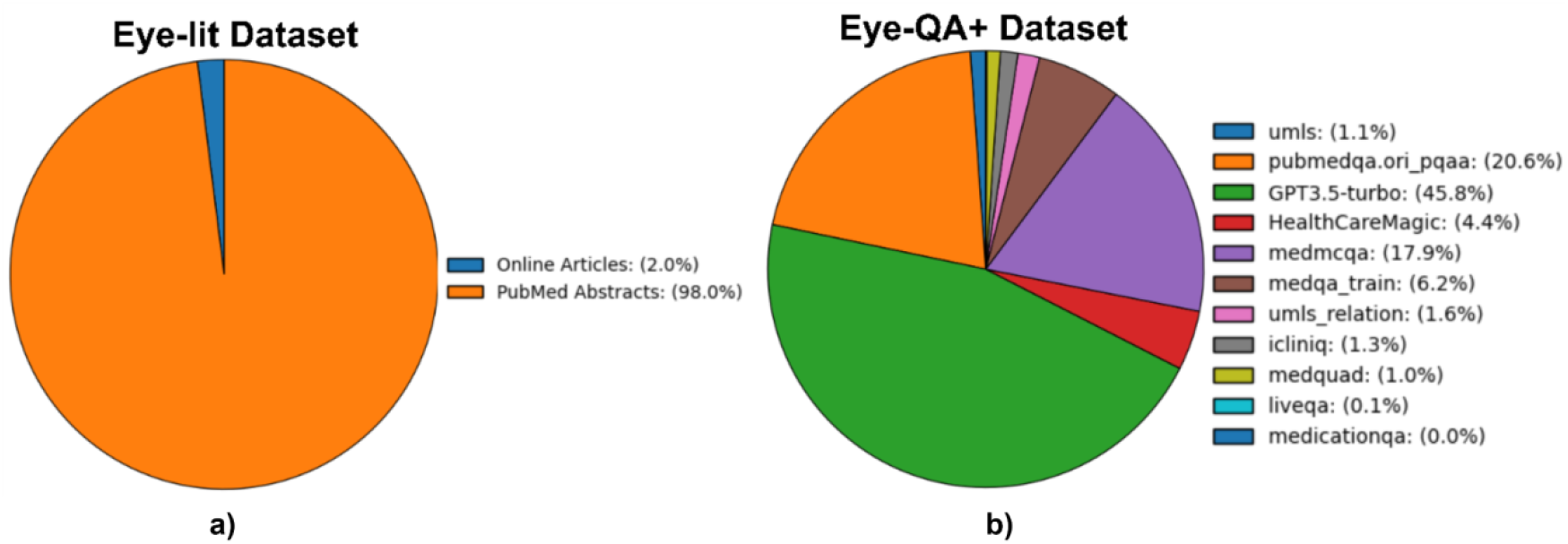
a) EYE-lit and b) EYE-QA+ datasets.

### 3.2 Model Training

We began with Llama 2-7B and trained it in two steps, capping sequence length at 512 tokens and using an effective batch size of 256. First, the model was pre-trained on the EYE-lit corpus, yielding **EYE-Llama_p**. We then fine-tuned that checkpoint on two successive Q&A collections: EYE-QA produced **EYE-Llama_qa**, and the larger EYE-QA+ set gave rise to **EYE-Llama_gqa**.

#### 3.2.1 Quantized Low-Rank Adaptation (QLoRA)

We adapted Llama 2-7B with **QLoRA**, a method that combines 4-bit (NF4) weight quantization with trainable low-rank adapters, so we could scale quickly without appreciable accuracy loss.^17,18^ Quantization slashes memory traffic while preserving the original weight distribution; the adapters provide the flexibility needed for new ophthalmic concepts.

Low-rank adapters supply the second half of the trick. Rather than updating every weight, we learn two slim matrices of rank *r* that capture tasks specific shifts. We set *r* = 16 after a small grid search showed it was the point where extra rank produced diminishing returns. The scaling term α = 64 modulates how much those new directions influence the frozen backbone. This configuration let us back-propagate through just twelve million parameters, barely 0.1 % of the model, yet still adapt to ophthalmic jargon and reasoning. This setup ran cleanly on two 49-GB GPUs: the pre-training stage logged 24 400 steps, followed by 300 and 500 fine-tuning steps for **EYE-Llama_qa** and **EYE-Llama_gqa**, respectively (Appendix D).

### 3.3 Evaluation

We evaluate the performance of our LLMs on multiple test sets, including a 780-question MedMCQA, an 800-question PubMedQA, a ∼1300 question patient/ophthalmologist synthetic QA pairs generated using GPT3.5, and a 20-question set designed by an ophthalmologist to qualitatively assess the LLMs, covering both MC and open-ended question formats.

#### 3.3.1 MCQA Evaluation

We evaluated the models on MC questions drawn from the MedMCQA and PubMedQA sets, reporting both accuracy and F1. For every response the model had to do more than pick a letter: it also had to defend that choice or explain why the remaining options fell short. Scoring was binary. A correct selection, paired with a coherent justification, earned a full point and signaled that the prompt was understood. Any wrong pick, along with answers that were missing, duplicated, or self-contradictory, received a zero, marking a lapse in comprehension or reasoning.

At inference time, we used the following prompt for the EYE-Llama_qa, EYE-Llama_gqa, Llama3, MedAlpaca, and GPT3.5/4 models:

*System prompt:* {*instruction*}

*User prompt:* {*question*}

And the following prompt for the ChatDoctor:

“{instruction}\n\nPatient: {question}\n\nChatDoctor: “

To perform inference with the Meditron model, we utilized their recommended prompts for both the PubMedQA and MedMCQA datasets. following prompt was used for PubMedQA:

*System prompt: “As an expert doctor in clinical science and medical knowledge, can you tell me if the following*

*statement is correct? Answer yes, no, or maybe.”*

*User prompt:* {*question*}

For the MedMCQA dataset, we utilized the following prompt:

*System prompt: “You are a medical doctor answering real-world medical entrance exam questions. Based on your*

*understanding of basic and clinical science, medical knowledge, and mechanisms underlying health*,

*disease, patient care, and modes of therapy, answer the following multiple-choice question. Select one*

*correct answer from A to D. Base your answer on the current and standard practices referenced in medical guidelines”*

*User prompt:* {*question*}

#### 3.3.2 Open-ended QA Evaluation

To assess how effectively our models respond to open-ended questions, we made use of two resources: the synthetic patient–ophthalmologist dialogue and the **20-Question**. The 20-Question dataset, available at huggingface.co/datasets/QIAIUNCC/EYE-TEST-2, was crafted by an ophthalmologist to probe clinical reasoning. Its prompts echo the questions a specialist would pose to an AI assistant, not the everyday concerns patients raise. Examples include, “What is the first-line therapy for primary open-angle glaucoma?” and “Which technique offers the best outcome for retinal detachment repair?” Two board-certified ophthalmologists independently reviewed every response, providing a consistent qualitative assessment.

For the open-ended Q&A datasets, we used the recommended prompt for ChatDoctor and Meditron models and the following prompt for the EYE-Llama_qa, EYE-Llama_gqa, and ChatGPT models:

*System prompt: “Given your profession as an ophthalmologist assistant, please provide a detailed and*

*comprehensive response to the question.”*

*User prompt:* {*question*}

## 4. Results

In this section, we provide an overview of our results as summarized in tables.

### 4.1 MedMCQA and PubMedQA datasets

**Table 2.** The table provides a performance comparison of various models.

| MedMCQA |  |  |
| --- | --- | --- |
| Model | Accuracy | F1-score |
| Llama 2-7b-chat | 0.33 | 0.19 |
| EYE-Llama_p | 0.34 | 0.21 |
| EYE-Llama_qa | 0.39 | 0.23 |
| EYE-Llama_gqa | 0.35 | 0.22 |
| ChatDoctor | 0.29 | 0.17 |
| ChatGPT (GPT3.5) | 0.55 | 0.37 |
| ChatGPT (GPT4) | 0.78 | 0.65 |
| Meditron-7B | 0.22 | 0.15 |
| Llama 3 | **0.90** | **0.80** |
| MedAlpaca-7B | 0.03 | 0.03 |
| PubMedQA |  |  |
| Model | Accuracy | F1-score |
| Llama 2-7b-chat | 0.9 | 0.45 |
| EYE-Llama_p | 0.74 | 0.34 |
| EYE-Llama_qa | **0.96** | **0.74** |
| EYE-Llama_gqa | 0.93 | 0.5 |
| ChatDoctor | 0.92 | 0.69 |
| ChatGPT (GPT3.5) | 0.93 | 0.52 |
| Meditron-7B | 0.76 | 0.34 |
| Llama 3 | 0.92 | 0.52 |

In the MedMCQA dataset, Llama 3 excels with superior accuracy and F1-score (Appendix E). Conversely, in the PubMedQA dataset, EYE-Llama_qa tops these metrics on the PubMedQA dataset. Note that the p-value for the PubmedQA and MedMCQA test sets is 0.05.

A noticeable performance decline is observed in the EYE-Llama_p and Meditron-7B models on the PubMedQA dataset. This can be attributed to the fact that the Llama 2 model, being chat-based, loses its task-specifisc knowledge (answering a question based on the provided context) after being pre-trained on unsupervised data. Nevertheless, the fine-tuned models (namely EYE-Llama_qa, and EYE-Llama_gqa) relearn this task during the fine-tuning phase, resulting in an improvement over its previous performance.

### 4.2 Open-ended Questions and Answers (QA)

#### 4.2.1 Synthetic dialogue dataset

The synthetic dialogue dataset, which includes ∼1300 QA, provided a broad range of patient inquiries and expert responses. The model’s performance on this dataset was quantified using the BERTScore, BARTScore and BLEU metrics.^19-21^

##### BERTScore, BARTScore and BLEU Metrics

We used BERTScore, BARTScore and BLEU Metrics to evaluate the model’s performance with the synthetic dialogue dataset. We used ground truth answers as the reference answer and the answers generated by each model as the candidate answer.

**Table 3.** Performance comparison of various models reveals the strong performance of the EYE-Llama_qa and EYE-Llama_gqa models across different metrics.

| synthetic dialogue |  |  |  |
| --- | --- | --- | --- |
| BERTScore |  |  |  |
| Model | Precision | Recall | F1 |
| Llama 2-7b-chat | 0.5194 | 0.6189 | 0.5630 |
| EYE-Llama_qa | 0.5392 | 0.6051 | 0.5702 |
| EYE-Llama_gqa | 0.5412 | 0.6056 | **0.5715** |
| ChatDoctor | **0.5933** | 0.5434 | 0.5648 |
| ChatGPT | 0.5295 | **0.6352** | 0.5757 |
| Meditron-7B | 0.5302 | 0.4874 | 0.5032 |
| Llama 3 | 0.5079 | 0.6229 | 0.5577 |
| BARTScore |  |  |  |
| Llama 2-7b-chat | **0.7054** |  |  |
| EYE-Llama_qa | 0.6322 |  |  |
| EYE-Llama_gqa | **0.7035** |  |  |
| ChatDoctor | 0.4553 |  |  |
| ChatGPT | 0.6689 |  |  |
| Meditron-7B | 0.6002 |  |  |
| Llama 3 | 0.6283 |  |  |
| BLEU score |  |  |  |
| Llama 2-7b-chat | 0.0154 |  |  |
| EYE-Llama_qa | 0.0212 |  |  |
| EYE-Llama_gqa | **0.0227** |  |  |
| ChatDoctor | 0.0170 |  |  |
| ChatGPT | 0.0193 |  |  |
| Meditron-7B | 0.0141 |  |  |
| Llama 3 | 0.0162 |  |  |

#### 4.2.2 Dataset of 20-Questions

Although automated scores exist, we used the 20-Questions dataset for a deliberately qualitative study focused on what practising ophthalmologists value in an answer. Two specialists created the questions and judged the model’s replies. For every response they considered six points:

- Alignment: Does the answer reflect current scientific consensus?
- Grasp: Did the model clearly understand the question?
- Relevance: Does it cite facts that genuinely help address the query?
- Irrelevance: Does it introduce incorrect or stray information?
- Harm: On a scale from 0 to 5, how much damage could the advice do?
- Overall quality: A holistic score from 0 to 10.

The first four items were graded with descriptive labels that map to numeric values: “Excellent/Yes” (1), “Mostly” (0.75), “Somewhat/Some” (0.5), “Mild” (0.35), “Minimal” (0.25), and “No” (0). The last two were scored directly on their respective scales (see Appendix B for full rubric).

**Table 4.**
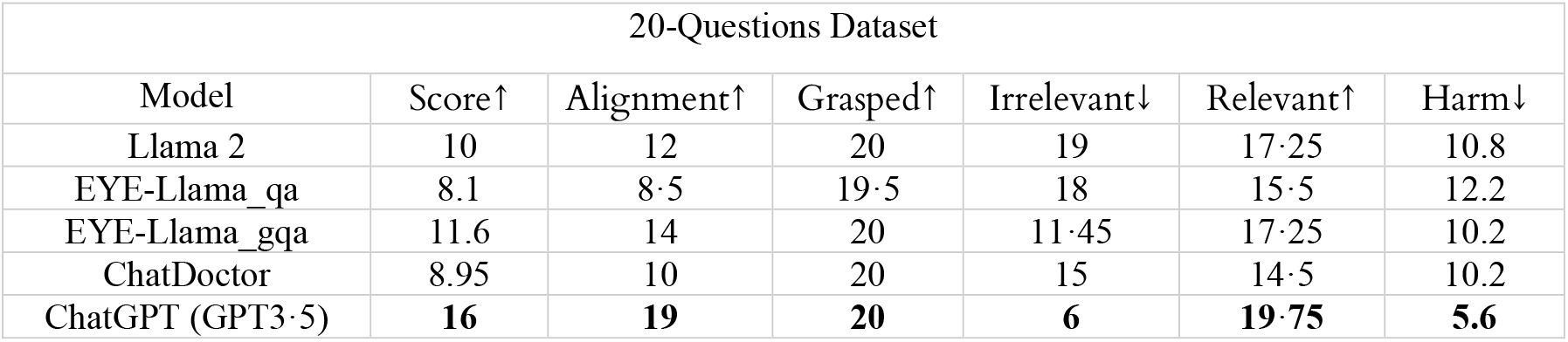
The table presents the scores (out of 20) assigned by ophthalmologists to the LLM responses for the 20-Question dataset.

## 5. Discussion

Our goal in this work was two-fold: first, to test whether a carefully curated ophthalmology corpus could lift the performance of a modest-sized backbone to the level of larger, general-purpose systems; and second, to show that memory-efficient fine-tuning methods make that jump feasible on mainstream hardware. Both objectives were met, though each leaves open avenues for refinement.

On MC benchmarks the pattern was consistent. After fine-tuning, EYE-Llama_qa rose to the top on PubMedQA, while EYE-Llama_gqa edged ahead on MedMCQA. In both cases the adapted models outperformed baseline Llama 2 and ChatDoctor and reached closer to the proprietary giants’ performance, ChatGPT and Llama 3. What changed was not model scale but data diet: an 18k sample of vetted ophthalmic QAs supplied the detail that generic corpora lack.

Open-ended evaluation told a complementary story. Automated scores (BLEU, BERTScore, BARTScore) placed EYE-Llama variants well ahead of baseline Llama 2 and roughly on par with ChatGPT, but the decisive evidence came from the 20-Question dataset. These twenty prompts, written and graded by practicing ophthalmologists, probe therapeutic strategy, procedural choice, and differential reasoning, domains where factual slip-ups can cause real-world harm. Across six qualitative axes, alignment, grasp, relevance, irrelevance, harm, and overall quality, EYE-Llama_gqa matched or exceeded ChatDoctor and untouched Llama 2, trailing only ChatGPT, whose 175 B parameters confer a clear advantage. In short, domain-tuned data plus targeted evaluation exposed gains that blanket metrics alone would have missed.

The performance jump rests squarely on the corpus we assembled. EYE-lit supplies 744 k PubMed abstracts, textbooks, and Wikipedia articles, giving the model exposure to canonical terminology and historical context. EYE-QA layers in 18 k supervised QAs, 1.5 k synthetic patient dialogues, and 15 k textbook-derived prompts, each vetted for coverage and accuracy. Finally, the 20-Question dataset offers a high-signal, low-noise probe of specialist reasoning. Together, these resources form a ladder: broad language at the base, focused QAs in the middle, and expert judgment at the top. Their public release should lower the barrier for future ophthalmic LLM work.

We fine-tuned with QLoRA 4-bit NF4 quantization plus rank-16 adapters (α = 64).^17,18^ Earlier studies noted that adapter schemes can lag full-matrix updates,^28^ yet here the memory drop came with negligible loss in accuracy. Twelve million learnable weights, 0.1 % of the model, proved enough to internalize ophthalmic jargon and reasoning. A direct ablation against full-precision fine-tuning remains future work, but the present results argue that adaptive quantization is a pragmatic choice when turnaround time matters.

Medical NLP has progressed rapidly since GPT-4 cleared the Ophthalmic Knowledge Assessment Program with an 81 % score, well above the 74 % human mean.^22–24^ Bernstein et al. later showed that eye-care chatbots produce answers so fluent that clinicians can separate humans from machines only 61 % of the time.^26^ Domain-tuned projects, Med-PaLM 2, BioGPT, Med-Alpaca, PMC_LLaMA, Ophtha-Llama2, OphGLM, reinforce the same lesson: feed a model the right data and it will specialize.^27,28,31,32^ Our contribution is to extend that pattern while making the underlying datasets open and traceable. Me-LLaMA is another recent effort that fine-tunes Llama 2 models on large-scale biomedical data, achieving strong performance on several medical NLP benchmarks.^12^

Several caveats warrant emphasis. First, genuinely labeled ophthalmic QA material is still thin on the ground: roughly one-third of EYE QA+ were generated with GPT 3.5 to patch obvious topical gaps. While each synthetic item was screened by an ophthalmologist for factual soundness, its prose inevitably mirrors the generator’s style and may imprint a bias that differs from authentic clinical dialogue. Second, our qualitative grading drew on only two expert reviewers; expanding the panel and measuring inter-rater agreement would provide a firmer statistical footing for future benchmarks.

From an ethical standpoint, every text source was public or fully de-identified, and citation links accompany each model answer to preserve traceability. Even so, large language models can “hallucinate” plausible sounding but incorrect statements; that risk is magnified in a clinical context. For that reason, EYE Llama is released strictly for research, education, and tool development purposes. Any move toward patient-facing deployment must include robust human oversight, external validation against real-world cases, bias audits on underserved populations, and compliance with emerging regulatory frameworks for AI in healthcare.

The next steps include expanding EYE-QA with real clinic notes, imaging captions, and longitudinal follow-ups; extending the 20-Question framework to cardiology, dermatology, and other subspecialties; and benchmarking QLoRA against full-precision updates under identical conditions. In parallel, bias audits and differential privacy filters will be crucial as more patient data enter the pipeline.

Our findings show that a 7 B-parameter model, armed with well-curated ophthalmic data and fine-tuned via QLoRA, can rival or surpass larger baselines on both structured and free-form tasks. By releasing EYE-lit, EYE-QA, and the 20-Question dataset, we hope to provide a platform on which others can build. With transparent data, open checkpoints, and clinician-in-the-loop evaluation, large language models may evolve into safe, reliable partners that lighten documentation, broaden access to specialist knowledge, and ultimately improve patient care.

## Supporting information

Supplemental Doc

## Resource Availability Lead contact

Requests for further information or materials should be directed to the lead contact, Minhaj Nur Alam.

## Materials availability

All code, trained model weights (EYE-Llama_p, EYE-Llama_qa, EYE-Llama_gqa), and datasets are publicly available. No new biological or chemical materials were generated in this study.

## Data and code availability

- EYE-lit (pre-training corpus): https://huggingface.co/datasets/QIAIUNCC/EYE-lit-complete (DOI: 10.57967/hf/5588)
- Synthetic dialogue QA: https://huggingface.co/datasets/QIAIUNCC/EYE-Synthetic-Dialogues (DOI: 10.57967/hf/5586)
- EYE-QA+ (expanded QA): https://huggingface.co/datasets/QIAIUNCC/EYE-QA-PLUS (DOI: 10.57967/hf/5226)
- 20-Questions test set: https://huggingface.co/datasets/QIAIUNCC/EYE-TEST-2 (DOI: 10.57967/hf/5228)
- MedMCQA test set: https://huggingface.co/datasets/QIAIUNCC/EYE-TEST-3 (DOI: 10.57967/hf/5229)
- PubMedQA test set: https://huggingface.co/datasets/QIAIUNCC/EYE-TEST-4 (DOI: 10.57967/hf/5230)
- Code: github.com/QIAIUNCC/EYE-Llama (DOI: 10.5281/zenodo.15272005)
- Eye-Llama_P model weights: https://huggingface.co/QIAIUNCC/EYE-Llama_p (DOI: 10.57967/hf/5223)
- Eye-Llama_qa model weights: https://huggingface.co/QIAIUNCC/EYE-Llama_qa (DOI: 10.57967/hf/5222)
- Eye-Llama_gqa model weights: https://huggingface.co/QIAIUNCC/EYE-Llama_gqa (DOI: 10.57967/hf/5224)

## Acknowledgments

This work was supported by the National Eye Institute grant R15EY035804 (MNA) and by the UNC Charlotte Faculty Research Grant (MNA)

## Author Contributions

T.H. led data curation, methodology development, and original draft writing, while S.G. performed visualization, contributed to methodological refinements, and assisted with manuscript writing. Supervision was provided by M.N.A., H.R., and F.H.; funding acquisition was carried out by M.N.A.; and model validation was performed by A.A., E.K., J.T.S., and T.L.

## STAR + METHODS

Detailed methods are provided in the online version of this paper and include the following:

## Key Resources Table

| REAGENT or RESOURCE | SOURCE | IDENTIFIER |
| --- | --- | --- |
| Deposited data |  |  |
| PubMed abstracts (ophthalmology keywords) | NCBI PubMed (Entrez API) | PMIDs available on request; search via https://pubmed.ncbi.nlm.nih.gov |
| Wikipedia ophthalmology articles | Wikipedia | https://en.wikipedia.org/wiki/Category:Ophthalmology |
| PubMedQA | Qiao Jin et al<sup>15</sup> | https://huggingface.co/datasets/llamafactory/PubMedQA |
| MedMCQA | Ankit Pal et al<sup>16</sup> | https://huggingface.co/datasets/openlifescienceai/medmcqa |
| ICliniQ and HealthCareMagic | Yunxiang Li et al<sup>9</sup> | https://huggingface.co/datasets/wangrongsheng/HealthCareMagic-100k-en |
| MedQuad | Asma Ben Abacha et al<sup>15</sup> | https://github.com/abachaa/MedQuAD/tree/master |
| Software and algorithms |  |  |
| Hugging Face Datasets | Hugging Face | https://github.com/huggingface/datasets |
| Hugging Face Transformers | Hugging Face | https://github.com/huggingface/transformers |
| QLoRA (Quantized LoRA) implementation | Tim Dettmers et al<sup>17</sup> | https://github.com/artidoro/qlora |

## Method Details

Data Collection and Preprocessing

1. **Keyword selection:** A curated list of ophthalmology terms (e.g., retina, glaucoma, macula) guided PubMed and QA dataset queries.
2. **PubMed abstracts:** Retrieved via BioPython Entrez in XML, raw text extracted, special characters removed, and irrelevant sentences (e.g., “in this paper”) filtered out.
3. **Textbooks & Wikipedia:** Downloaded PDFs/HTML, segmented into ∼512-token chunks to match abstract lengths.
4. **Supervised QA filtering:** Subsets of PubMedQA, MedMCQA, PMC_LLaMA, Icliniq, HealthCareMagic, and MedQuad selected by keyword.
5. **Synthetic dialogues:** Prompted GPT-3.5 to produce ∼1,500 patient-style Q&A pairs for training (Appendix A1); ophthalmologists reviewed and curated these.
6. **Training dataset assembly:**

∘ **EYE-lit:** ∼744 k abstracts + textbook & Wikipedia chunks
∘ **EYE-QA:** 18 k supervised QAs + 1.5 k synthetic dialogues
∘ **EYE-QA+:** EYE-QA + 15 k additional GPT-3.5–generated QAs (Appendix A2)

## Model Training

1. **Base model:** Llama 2-7b-chat loaded with Hugging Face Transformers.
2. **Pre-training (EYE-Llama_p):**
  ∘ Sequence length: 512 tokens
  ∘ Batch size (effective): 256
  ∘ Steps: 24,400
  ∘ QLoRA quantization + LoRA adapters (r = 16, α = 64) enabled fine-tuning on two NVIDIA A40 GPUs.
3. **Fine-tuning:**
  ∘ **EYE-Llama_qa:** Trained on EYE-QA for 300 steps.
  ∘ **EYE-Llama_gqa:** Trained on EYE-QA+ for 500 steps.
4. **Optimization:**
  ∘ Learning rate scheduler: linear warmup + decay
  ∘ Gradient accumulation to achieve effective batch size when GPU memory limited.

## Evaluation Protocols

- **MCQA tests:** MedMCQA (780 questions) and PubMedQA (800 questions).
- **Open-ended tests:**
  ∘ **Synthetic dialogue:** ∼1,300 QAs with GPT-3.5 reference answers.
  ∘ **20-Questions:** 20 clinician-designed queries; responses graded qualitatively by two ophthalmologists.
- **Metrics:** Accuracy and F1 for MCQA; BERTScore, BARTScore, BLEU for synthetic dialogues; custom multi-axis scoring (alignment, grasp, relevance, harm, overall) for 20-Questions.

## Quantification and Statistical Analysis

Quantification and Statistical Analysis

- **Accuracy**: Proportion of exactly correct MC answers (justification ignored).
- **F1 score**: Harmonic mean of precision and recall over MC options.
- **BERTScore**: Computed with Hugging Face library (Cosine similarity on contextual embeddings).
- **BARTScore**: Evaluates generated text by computing the likelihood of reconstructing the reference text using a pre-trained BART model, effectively treating evaluation as a text generation task.
- **BLEU**: A metric that assesses the quality of machine-translated text by comparing it to one or more reference translations, focusing on the precision of n-gram matches.

## Qualitative scoring

- **Alignment, grasp, relevance, irrelevance**: Rated on a 0 –1 scale (Excellent = 1, Mostly = 0.75, Somewhat = 0.5, Mild = 0.35, Minimal = 0.25, No = 0).
- **Harm**: On a 0 – 5 scale, rescaled to 0 – 20.
- **Overall score**: On a 0 –10 scale, rescaled to 0 – 20.

**Software**: All analyses performed in Python 3.10 with NumPy, SciPy, and Hugging Face Evaluate.

## Supplemental Information

Supplemental information can be found in EYE Llama Supplementary file.

