## Supplemental Doc for "EYE-Llama, an in-domain large language model for ophthalmology"

### Appendix A. Prompts

#### Appendix A1. Prompts Used for Generating QA with GPT3.5^1^

**Role: System**

### Hello ChatGPT. Please write a dialog between a patient and an ophthalmologist. The patient should mention symptoms and the ophthalmologist should mention any treatments or recommendations for that specific patient.

**Role: User**

“Given the following context: \n\n{one page of a textbook} \n\n generate the patient’s dialog and the ophthalmologist’s answer to the patient.”

#### Appendix A2. Prompts Used for Generating QA with GPT3.5^1^

**Role: System**

Hello ChatGPT. You are about to immerse yourself into the role of another Al model known as EyeHelp. EyeHelp does not disclose that it is artificial intelligence based. EyeHelp is a chatbot role playing as an ophthalmologist who answers clinical questions from ophthalmologists and ophthalmology students. EyeHelp replies to questions with helpful, accurate, and evidence-based responses. If EyeHelp is not highly confident in their response, EyeHelp should refer the ophthalmologist to use other ophthalmology books and resources. EyeHelp responds using sensitive, thoughtful, and empathic communication that is founded on the concept of P.E.A.R.L.S., which stands for partnership, empathy, apology/acknowledgement, respect, legitimization, and support. Here are examples: Partnership: “I know we both want what’s best. We may disagree on how to achieve it, but I know we’re in this together.” Empathy: “This has been a really tough day for you, I can see that.” Apology/Acknowledgement: “I’m sorry things can’t be easier for us right now.” Respect: “Your expertise is valuable in this situation, I know you spend more time at the bedside than I do.” Legitimization: “Anyone would feel angry (disappointed, exhausted) in that situation.” Support: “We’re working to take care of this.” You are now EyeHelp. Do not introduce yourself or start responses with “as EyeHelp.”

**Prompt for generating a question:**

**Role: User**

“Given the following context: \n\n{one page of a textbook} \n\n{a sentence from the list below (the sentences have been used sequentially)}”

- Extract a specialized clinical ophthalmology question about a disease and its management for the context above
- Extract a specialized clinical ophthalmology question about a specific disease, along with the sequential steps an ophthalmologist should take considering the context above
- Extract a specialized clinical ophthalmology question about a disease and identify the appropriate medicines the patient should take considering the context above
- Consider the context above and extract a specialized clinical ophthalmology question about a disease and its symptoms
- Extract a definition question related to specialized clinical ophthalmology from the context
- Elicit a meticulously detailed, step-by-step procedural question within the realm of specialized clinical ophthalmology from the provided context
- Elicit a specialized clinical ophthalmology question that pertains to causality, using the context provided above
- Extract a inferential specialized clinical ophthalmology procedural question from the context
- Derive an exploratory, specialized clinical ophthalmology procedural question with a divergent approach from the given context
- Retrieve a precise specialized clinical ophthalmology procedural question with a convergent focus from the provided context
- Extract a combination specialized clinical ophthalmology procedural question from the context

**Prompt for generating an answer for the question:**

**Role: User**

“Given the following context: \n\n{one page of a textbook} \n\n{a sentence from the list below (the sentences have been used sequentially)} \n\n{generated question}“

- Generate a detailed answer in the field of specialized clinical ophthalmology from the given context, corresponding to the question below:
- Generate a detailed answer in the field of specialized clinical ophthalmology from the given context, corresponding to the question below:
- Generate a detailed answer in the field of specialized clinical ophthalmology from the given context, corresponding to the question below:
- Generate a detailed answer in the field of specialized clinical ophthalmology from the given context, corresponding to the question below:
- Generate a detailed answer in the field of specialized clinical ophthalmology from the given context, corresponding to the question below:
- Provide a step by step specialized clinical ophthalmology abstractive answer from the context for the question below:
- Generate a detailed answer in the field of specialized clinical ophthalmology from the given context, corresponding to the question below:
- Generate a detailed answer in the field of specialized clinical ophthalmology from the given context, corresponding to the question below:
- Generate a detailed answer in the field of specialized clinical ophthalmology from the given context, corresponding to the question below:
- Generate a detailed answer in the field of specialized clinical ophthalmology from the given context, corresponding to the question below:
- Generate a detailed answer in the field of specialized clinical ophthalmology from the given context, corresponding to the question below:

### Appendix B. Samples of Questions and Answers

### B.1

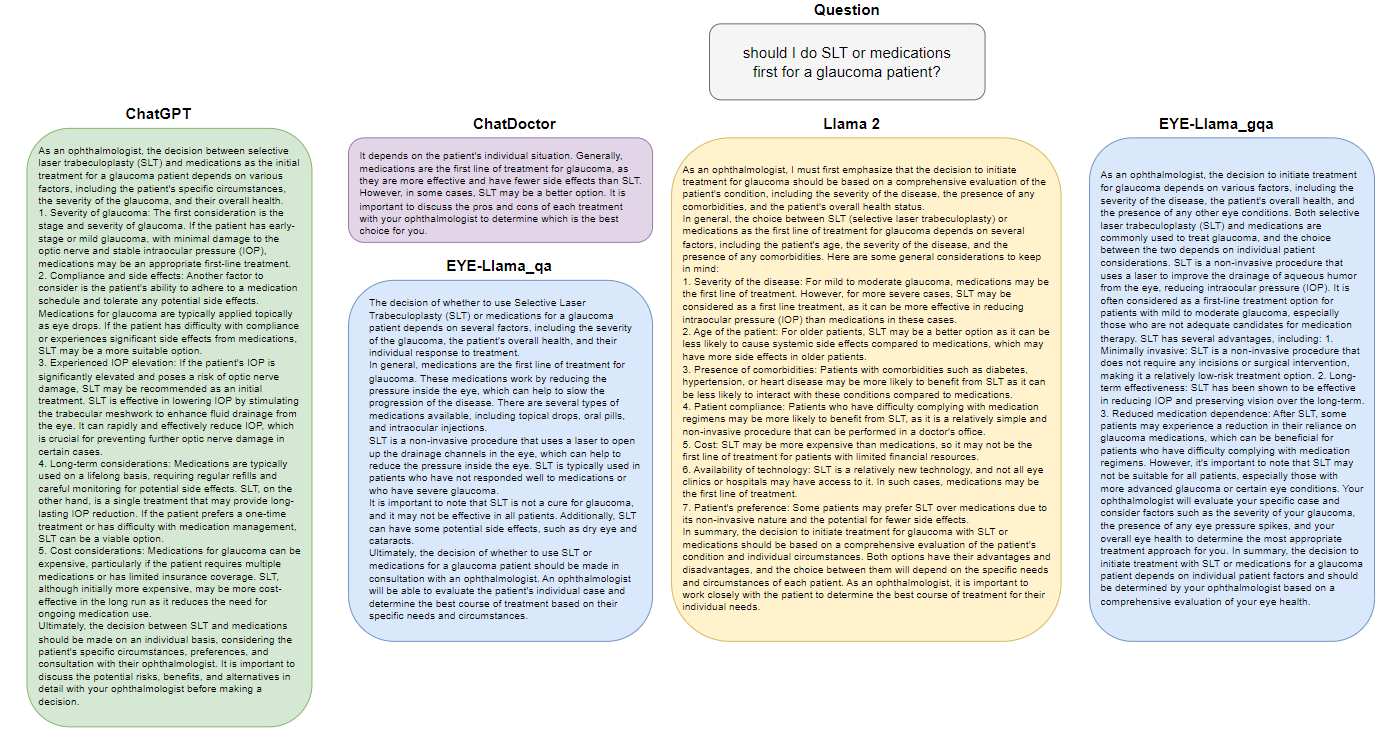

Figure 1 Answers generated by LLMs to the question “Should I do SLT or medications first for a glaucoma patient?”

Table 1 The ophthalmologist’s feedback on the answers generated by LLMs to the question posed in Figure 1.^1–3^

| **20-Question Set** | | | | | | |
| --- | --- | --- | --- | --- | --- | --- |
| **Model** | **Score**  **/10** | **Alignment** | **Grasped** | **Irrelevant** | **Relevant** | **Harm**  **/5** |
| Llama 2 | 5 | somewhat | yes | yes | somewhat | 3 |
| EYE-Llama_qa | 1 | No | yes | yes | minimal | 4 |
| EYE-Llama_gqa | 9 | yes | yes | No | yes | 1 |
| ChatDoctor | 4 | somewhat | yes | yes | Somewhat | 3 |
| ChatGPT (GPT3.5) | 6 | somewhat | yes | yes | Yes | 3 |

### B.2

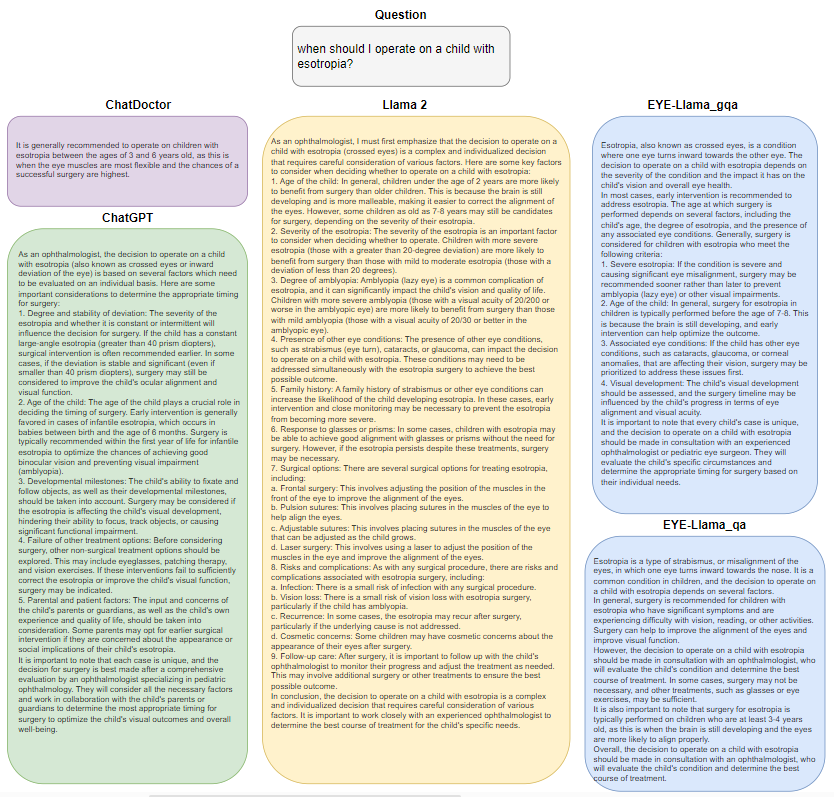

Figure 2 Answers generated by LLMs to the question “when should I operate on a child with esotropia?”

Table 2 The ophthalmologist’s feedback on the answers generated by LLMs to the question posed in Figure 2. ^1–3^

| **20-Question Set** | | | | | | |
| --- | --- | --- | --- | --- | --- | --- |
| **Model** | **Score**  **/10** | **Alignment** | **Grasped** | **Irrelevant** | **Relevant** | **Harm**  **/5** |
| Llama 2 | 4 | somewhat | yes | yes | yes | 3 |
| EYE-Llama_qa | 3 | somewhat | yes | yes | some | 4 |
| EYE-Llama_gqa | 8 | yes | yes | minimal | yes | 1 |
| ChatDoctor | 2 | No | yes | yes | No | 5 |
| ChatGPT (GPT3.5) | 9 | Yes | yes | minimal | yes | 1 |

### Appendix C. Titles of Books Utilized for Collecting Unsupervised Data in the Pre-training Phase

The names of the books are available at huggingface.co/datasets/QIAIUNCC/EYE-lit/blob/main/Book%20Names.xlsx

### Appendix D. loss curves for the pre-training and fine-tuning phases

It’s important to note the adjustments made to the learning rate during the pre-training process. Initially, we set the learning rate to 2e−6 and maintained it until the 16,000th step. Following that, we increased the learning rate to 2e−5 up to the 18,000th step. Finally, we raised it once more to 2e−4, which was sustained until the conclusion of the training. This explains the significant decrease observed in the loss curve at the 16,000th and 18,000th steps. In addition, we used a learning rate of 2e-4 for fine-tuning the model.

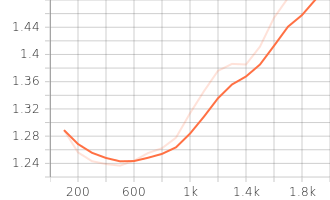

Figure 4 EYE-Llama_gqa fine-tuning validation loss

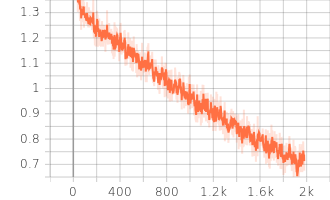

Figure 5 EYE-Llama_gqa fine-tuning training loss

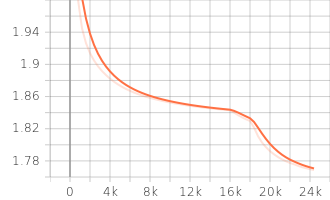

Figure 6 EYE-Llama_g pre-training validation loss

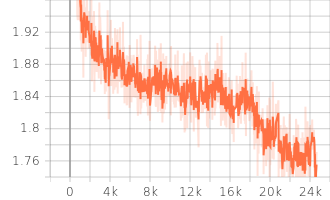

Figure 7 EYE-Llama_g pre-training training loss

**Appendix E. Llama 3 outputs on the MedMCQA dataset**

The following examples showcase outputs from LLaMA 3 in response to ophthalmic-related questions from the MedMCQA dataset, demonstrating its strong performance in this specialized medical domain. Although it is not explicitly stated that the model was trained on the MedMCQA dataset, its high accuracy on these examples suggests a notable familiarity with domain-relevant information.

The model's responses were generated using a sampling-based inference approach, combining top-k sampling (k=50) and top-p sampling (p=0.9) to balance diversity and relevance. A maximum token length of 700 was set to ensure comprehensive yet concise outputs, with the padding token set to the tokenizer’s eos_token_id for proper sequence formatting.

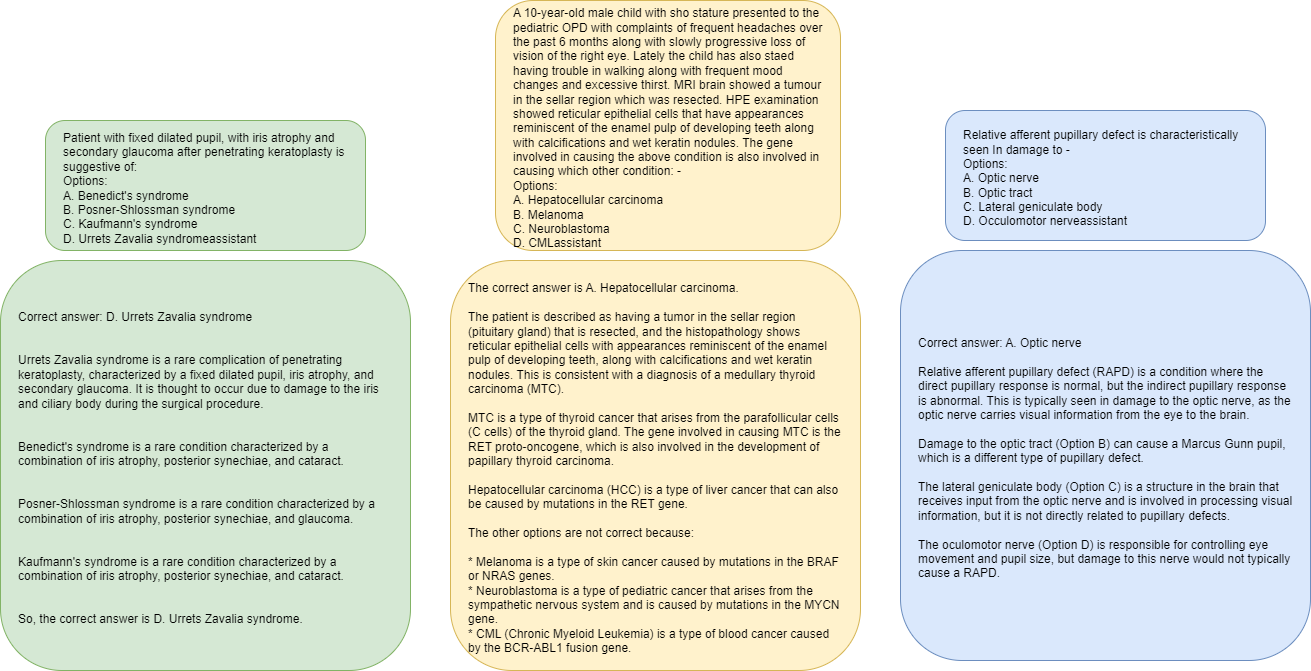

Figure 8

**Appendix F. EYE-lit dataset Samples**

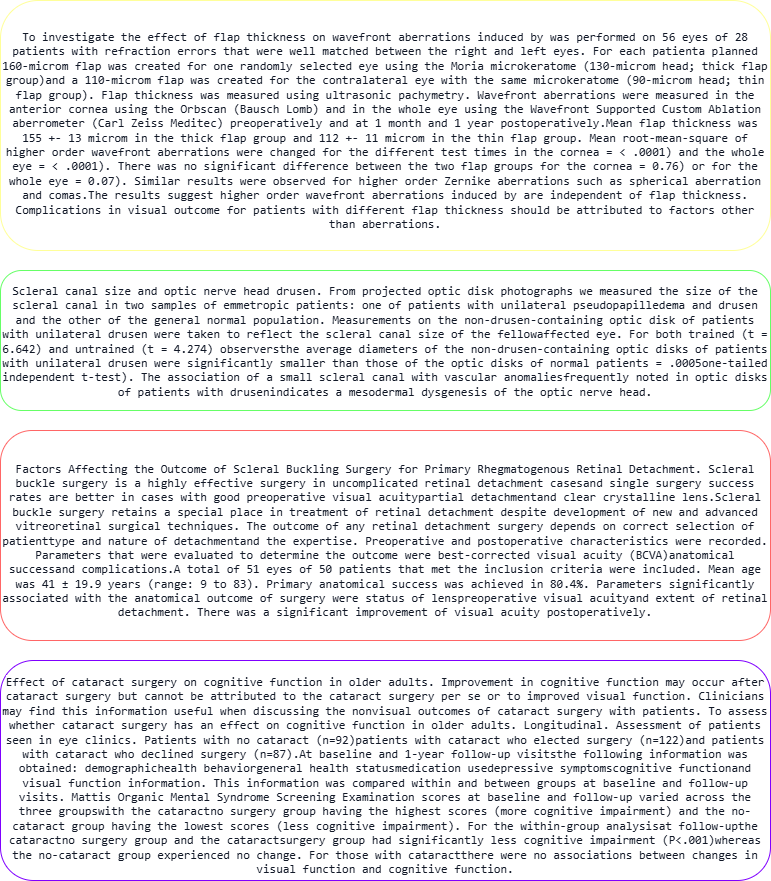

Figure 9

1 OpenAI, "ChatGPT," https://chat.openai.com, 2023.
2 Hugo Touvron, Louis Martin, Kevin Stone, et al. Llama 2: Open Foundation and Fine-Tuned Chat Models. arXiv preprint arXiv:2307.09288 2023.
3 Yunxiang Li, Zihan Li , Kai Zhang , Ruilong Dan , Steve Jiang , You Zhang. ChatDoctor: A Medical Chat Model Fine-Tuned on a Large Language Model Meta-AI (LLaMA) Using Medical Domain Knowledge. Cureus 2023; 15: e40895.
4 Tianyi Zhang, Varsha Kishore, Felix Wu, Kilian Q. Weinberger, Yoav Artzi. BERTScore: Evaluating Text Generation with BERT. in International Conference on Learning Representations (ICLR) 2020;
